# Tracking spatial patterns and nocturnal arousal in an undisturbed natural population of the pulse-type weakly electric fish *Gymnotus omarorum*

**DOI:** 10.1101/2024.06.29.600875

**Authors:** Adriana Migliaro, Federico Pedraja, Stefan Mucha, Jan Benda, Ana Silva

**Author notes:** Lead contact: Ana Silva. These authors contributed equally to this work and shared co-first authorship (Adriana Migliaro, Federico Pedraja).

## Abstract

Assessing animals’ locomotor and activity-rest patterns in natural populations is challenging. It requires individual identification and behavioral tracking in sometimes complex and inaccessible environments. Weakly electric fish are advantageous models for remote monitoring due to their continuous emission of electric signals (EODs). *Gymnotus omarorum* is a South American freshwater pulse-type weakly electric fish. Previous manual recordings of restrained individuals in the wild showed a spatial distribution compatible with territoriality and a nocturnal increase in EOD rate interpreted as arousal. This interdisciplinary study presents the development of low-cost amplifiers for remote EOD recordings and the refinement of tracking algorithms that provide individual recognition of *Gymnotus omarorum* in the wild. We describe natural daily spacing patterns of undisturbed individuals that are compatible with territoriality, although heterogeneous across sampling sites, and confirm that all resident fish showed a robust nocturnal increase of EOD rate likely associated with daily variations of water temperature.

**HIGHLIGHTS:**

1. Successful remote individual tracking of wild pulse type weakly electric fish
2. *G. omarorum* spacing patterns are compatible with known nocturnality and territoriality
3. Residents keep their diurnal resting sites and move within small areas during the night
4. The robust nocturnal electric arousal of residents is linked to water temperature peak

## INTRODUCTION

Evaluating locomotor and activity-rest patterns in natural populations of animals has been and still is a big challenge^1^. Monitoring animals in the wild requires feedback between high-precision measurements of the behaviors of undisturbed animals, as well as mathematical and computational algorithms that seek to recreate these behaviors from the measurements. Recent advances in field-deployable tracking technologies (e.g. stationary imaging techniques, bio-loggers and remote sensing) present new opportunities for conducting field-based studies at ecologically meaningful spatio-temporal scales^2^. However, these techniques have their own limitations. It remains challenging to identify the proper site of remote sensing to get reliable data, or to monitor small animals that cannot carry individual loggers, or to track animals that live in inaccessible or complex environments for video recordings. On top of that, tracking requires individual identification and therefore a reliable individual-specific trait to act as a biometric cue, which is an additional challenge for the resolution of the available recording devices^3–5^.

Freshwater weakly electric fish, that have evolved independently in South America and Africa, generate weak electric fields to locate and identify nearby objects (electrolocation) and to communicate with each other (electrocommunication)^6,7^. Electrocytes densely packed into the peripheral electric organ fire in synchrony, driven by spinal electromotor neurons that receive input from a hindbrain command circuit, to generate an external electric field^8,9^. Electric fish are advantageous models for remote monitoring because they continuously broadcast electric information, which can be easily recorded remotely without disturbing natural populations^10^.

Furthermore, to serve as communication signals, these electric organ discharges (EODs) carry information about species identity, sex, social status, and motivational state^8^. Wave-type weakly electric fish generate continuous, highly periodic EODs, whose high frequencies (200-1000 Hz) and waveforms are species- and individual-specific. Pulse-type weakly electric fish emit short (typically less than 2 ms) stereotyped pulses with longer periods of silence, whose rate is lower (<100 Hz) and highly variable in mormyriforms but quite regular in gymnotiforms^11^. Therefore, EODs can also be used as a cue for individual recognition, although several species (especially pulse-type electric fish) show dramatic plastic changes in their EOD rates and waveforms, which can jeopardize individual recognition, particularly in natural scenarios^12,13^. Recent studies have succeeded in the remote tracking of wave-type electric fish in the wild based on the individual-specific EOD frequency^3,10,13,14^. However, given that the EOD rate of pulse-type electric fish is highly variable and context dependent, individual monitoring has only been achieved so far with the help of video tracking and machine learning, methods that are not feasible to implement in the wild^15,16^.

*Gymnotus omarorum* is a pulse-type weakly electric fish widely distributed in the southern boundary of gymnotiform distribution in South America^17,18^. It has been extensively studied as a model system for the understanding of the anatomo-functional principles of active electroreception^19^ and of the neuroendocrine mechanisms underlying intra- and inter-sexual year-round territorial aggression^20,21^. Ecological data obtained from manual electric census during the resting phase of *G. omarorum* in the wild showed a spatial distribution compatible with territoriality^22^. Individual *G. omarorum* were distributed evenly, and not aggregated with other conspecifics under the core area of the floating mats in constant darkness but were absent at the edge of the plant cover near the open water^22^.

Although fish were found in the same spot on successive days of the electric census, this manual procedure did not allow to track individuals during their active night phase or verifying site fidelity, which require individual recognition^23,24^. Previous ecological data have also shown that *G. omarorum* is a nocturnal fish that increases both its exploratory behavior and its electric behavior during the night^25^. The nocturnal arousal of weakly electric fish can be observed by the increase in EOD basal rate that functions as an alert mechanism that improves electroreceptive resolution^26–29^. Manual recordings of the EODs from identified individuals *G.omaroum* with restricted movements in both laboratory^29^ and natural settings^25^ showed a robust nocturnal increase of EOD basal rate, which persisted in total darkness and in motionless fish. Therefore, the environmental light cycle seems not to be a relevant trigger for the nocturnal electric arousal of *G. omarorum,* providing the opportunity to evaluate the alternative synchronizing role of the water temperature daily cycle^25^. However, the nocturnal electric arousal and its temporal relation to the daily temperature cycle have never been recorded in undisturbed animals in their natural shelters. All these biologically relevant questions could be answered by the automatization of remote EOD recordings of undisturbed individuals of *G. omarorum* in their natural habitat, which has faced several methodological barriers so far.

In this interdisciplinary study, we present the development of an updated version of low-cost amplifiers that allow remote recordings of the electric behavior of *G. omarorum* in the wild, as well as a tracking algorithm to provide the first individual recognition of fish within a natural population of pulse-type weakly electric fish. In addition, the application of this methodology to a well-known natural, undisturbed population of *G. omarorum* allowed us to shed light on three of the open questions posed by previous evaluations of this same population that remain unanswered because of the limitations of the non-automatized procedures used so far^22,25^. First, is there a differential spatial distribution of fish across the littoral zone of the lake homogeneously covered by aquatic plants? Second, is there a day-night change in these spatial patterns? Third, do undisturbed natural individuals of *G. omarorum* display the same nocturnal electric arousal as previously reported in non-freely moving animals?

## RESULTS

The electric behavior of undisturbed individuals of the pulse-type gymnotiform, *Gymnotus omarorum,* was recorded in their natural habitat (Laguna de los Cisnes; 34° 480 S, 55° 180 W), a 205 ha freshwater lake located in Maldonado, Uruguay, around the fall (March) and spring (October) equinox in 2023 (Figure 1). *Gymnotus omarorum* is the only gymnotiform species that occurs in this site^18^ and that rests during daytime underneath the roots of a thick strip of free-floating plants that cover the littoral area of this lake (Figure 1B). Following Zubizarreta et al.^3^, who found that individuals of *G. omarorum* preferred to rest at the core of this homogeneous littoral area but not at the edge, we placed one 8-channel recording device on the core sampling area and a second one on the edge, respectively (Figure 1B). We obtained simultaneous continuous recordings from both sampling areas over a 1000-minute period (approximately 17 hours), covering the transition from day to night, the entire night, and the transition from night to day. Recordings were taken on three different nights, each night at different positions within the core and the edge sampling areas. The eight recording electrodes of the recording devices were arranged in a grid structure (120 x 80 cm, with electrodes spaced at 40 cm; Figure 1C). EODs of *G. omaroum* individuals were recorded onto microSD cards using low-cost, battery-operated Teensy microcontrollers and custom-made multi-channel amplifiers^30^; Figure 1D-E; Figure S1).

**Figure 1.**
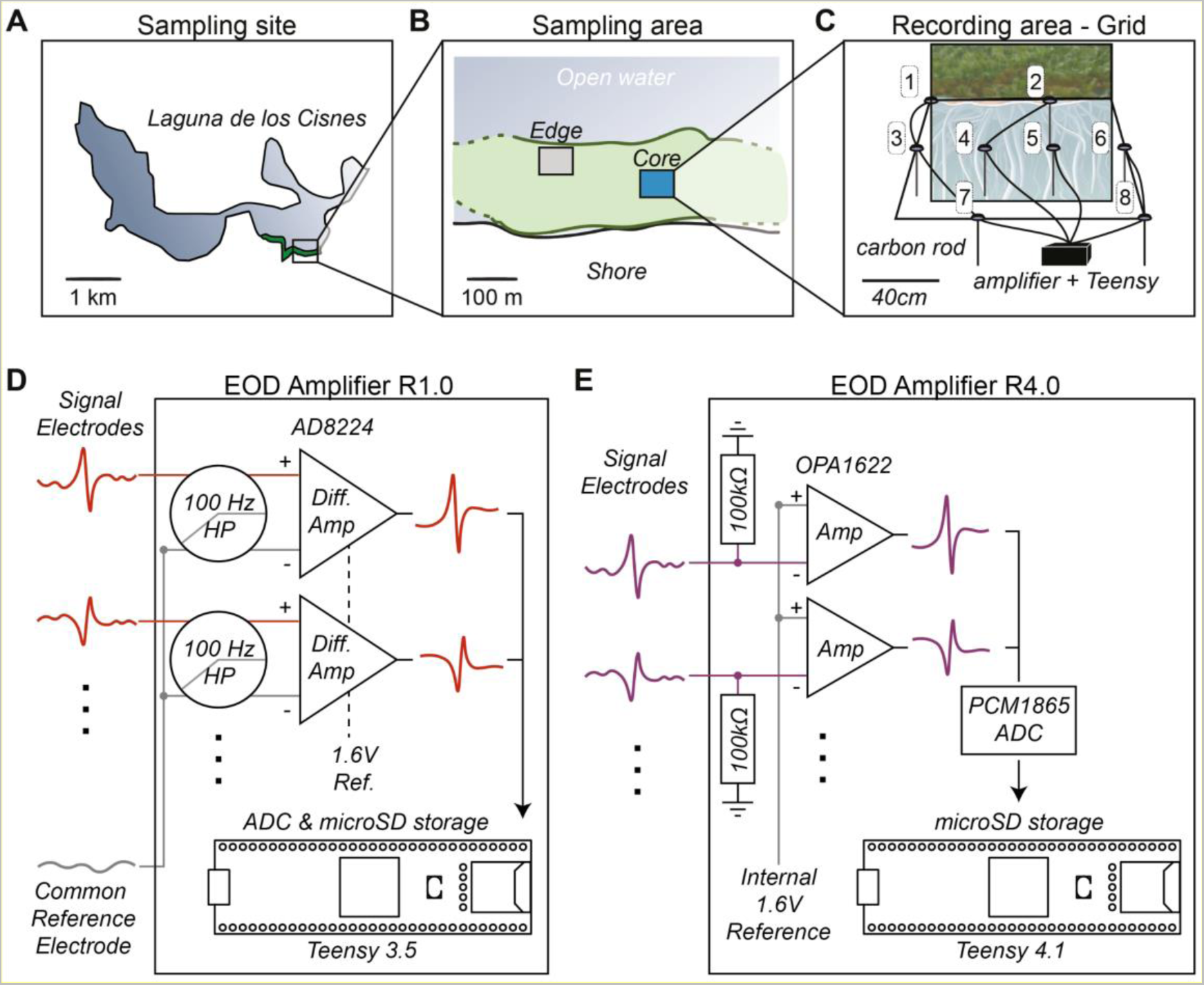
Representation of sampling sites and the automatized recording system. (A) The study site is located in Maldonado, Uruguay, in Laguna de los Cisnes. (B) Water hyacinths along the shore of the lake create an extensive floating mat that constitutes the sampling area. The recording devices were placed at both the core (blue rectangle) and the edge (grey rectangle). (C) Schematic representation of the recording device: An array of an eight-electrode grid system (120 x 80 cm, with electrodes spaced at 40 cm) was positioned on top of the floating mats, with the carbon electrodes (40 cm) submerged underwater through the plant roots. (D) Schematic of the R1.0 EOD amplifier. Signals were high-pass filtered (HP, 100 Hz cutoff frequency) and differentially amplified with a gain of 30 against a common reference electrode that was placed at a distance of 50 cm to the recording electrodes in the water. Amplified signals were digitized with a sampling rate of 20 kHz and stored in 16-bit wave files on a microSD card using a Teensy 3.5 with onboard analog-digital converter (ADC). (E) Schematic of the R4.0 EOD amplifier. Signals were coupled to battery ground (GND) via 100 kOhm resistors and amplified against an internal 1.6V reference with a gain of 10. Note that this setup does not use an external reference electrode. The signals were further amplified (gain of 10) and digitized with a sampling rate of 48kHz using a PCM1865 ADC and stored on a microSD card using a Teensy 4.1.

A two-step algorithm was used to extract the positions and identities of freely moving individuals in the wild from the EOD recordings as shown in Figure 2. First, the algorithm tracked the spatial position of each fish in relation to the grid geometry using the relative EOD amplitude across grid electrodes based on weighted spatial averages as previously described^14^ (Figure 2B). Second, the algorithm assigned an identity to each fish through k-means clustering of spatial position previously obtained in the first step and applied a merging criterion to contiguous clusters that exhibited similar EOD rates (Figure 2C). To illustrate the discrimination power of this tracking algorithm, Figure 2A shows 2 seconds of the raw recordings from the first 100 min recording of one of the core sampling sites (Core 1 in Figure 3) in which 5 out of the 8 electrodes of the grid system captured EODs. The application of the tracking algorithms allowed us to identify the presence and position of 2 distinct individuals of *G. omarorum* (magenta and cyan fish) underneath the grid system (Figure 2B) as well as the transition of one of them (the cyan fish from close to electrode #6 to close to electrode #2; Figure 2C) during this recording period. Once each detected fish was identified individually, we calculated their EOD rates (as the mean value of stable 1-min EOD recordings) and analyzed its variations throughout the day-night-day transitions. Figure 2B shows the values of the EOD rate of each identified individual during this recording period, which was slightly higher in the cyan fish (33.5 Hz) than in the magenta fish (31.5 Hz).

**Figure 2.**
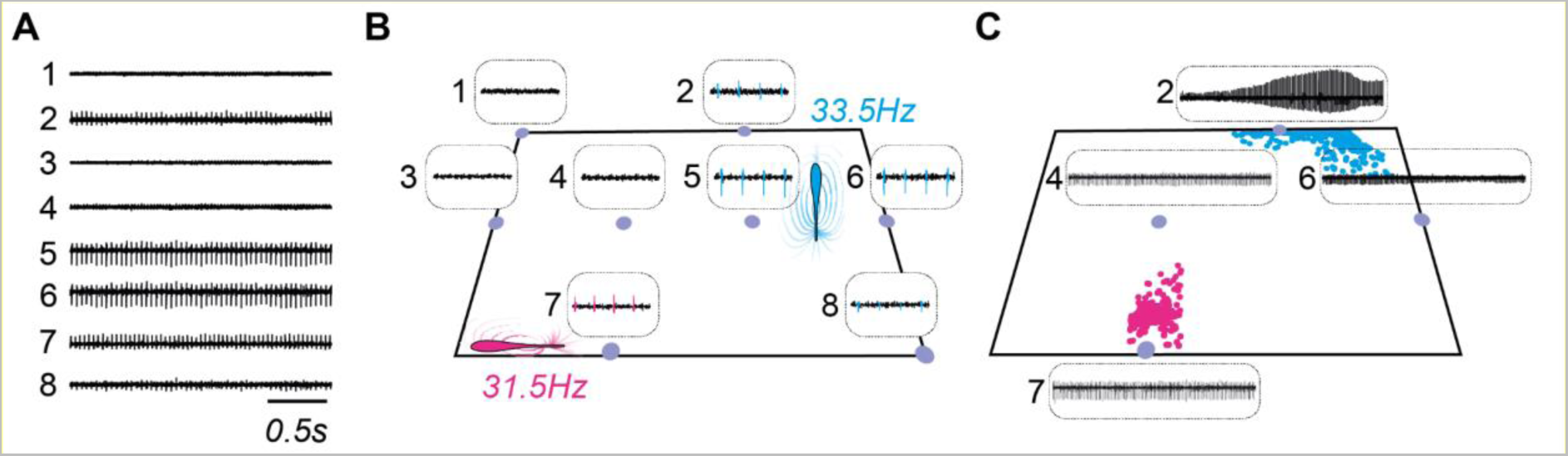
EOD recordings in precise positions allow for individual identification and fish position estimation. (A) Recorded traces from the 8-channels grid showing EODs for 2 s from the first 100 min recording of Core 1 sampling site. (B) Example of fish position and identity based on EOD signals obtained from the different electrodes. In this case, two fish (magenta and cyan) were identified, and a subsequent calculation of the EOD rate was performed. (C) Fish position over time from the example in (B). The change in EOD amplitude reflects a change in fish position. In this case, a transition between different parts of the grid can be observed for one fish (cyan).

**Figure 3.**
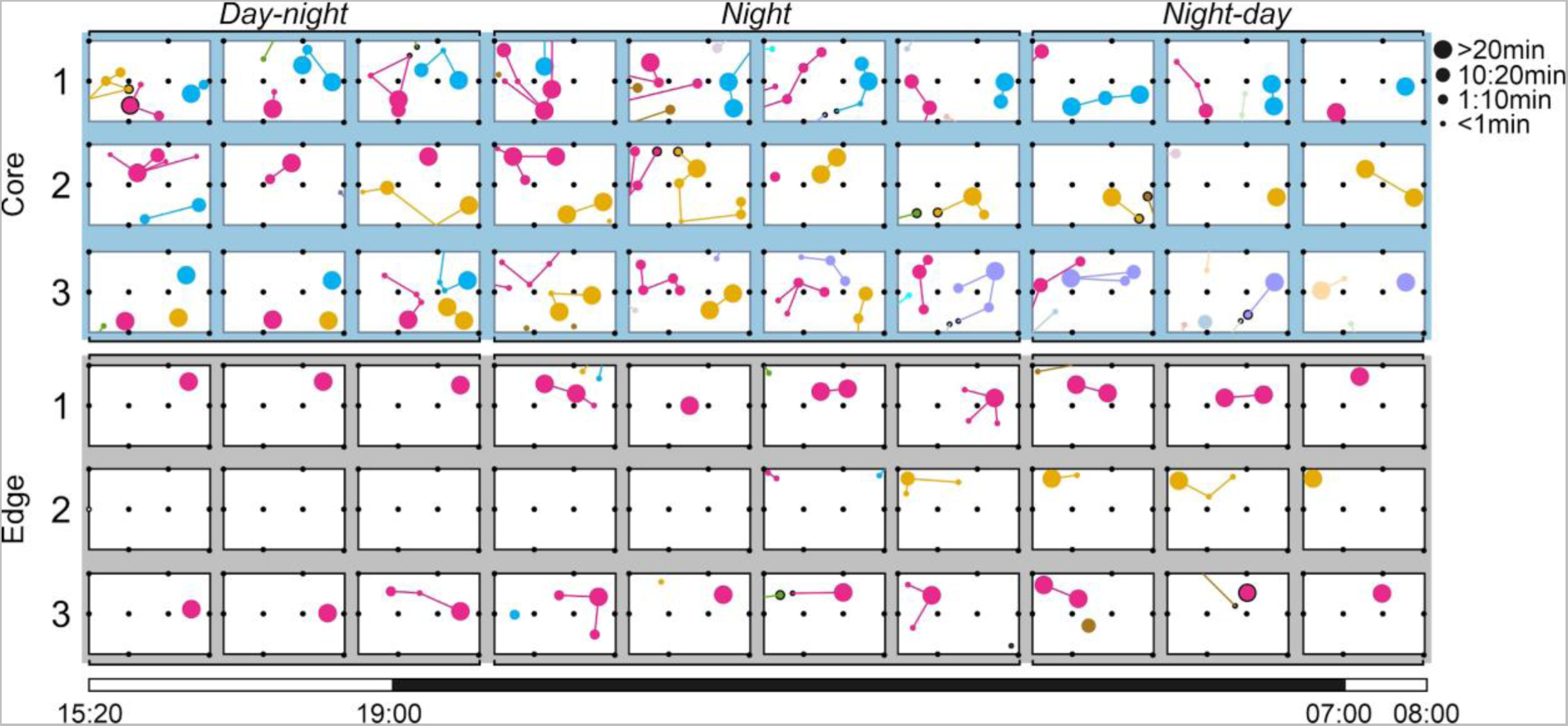
Distinctive spatial patterns between core and edge sampling areas. Data obtained from the monitoring of 6 sampling sites, with three situated at the core of the littoral area (upper rows, light blue) and three at the edge of the littoral area (lower rows, grey), recorded from 15:20 to 08:00 in 10 100-minute windows. Each tracked individual is represented by a dot, distinguished by its distinctive color (the first identified fish was marked in magenta, the second in cyan, the third in yellow, and so on), enabling tracking of its position over the 1000-minute recording time. The size of the dots in each 100-minute window reflects the duration for which each fish was recorded in the tracked position, while transitions between positions are indicated by connecting lines. Fish were categorized as residents when they were tracked for more than 60 minutes in the day-night and/or night-day transitions: C1R1 (magenta fish in row Core 1), C1R2 (cyan fish in row Core 1), C2R1 (magenta fish in row Core 2), C2R2 (yellow fish in row Core 2), C3R1 (magenta fish in row Core 3), C3R2 (cyan fish in row Core 3), C3R3 (yellow fish in row Core 3), C3 R4 (purple fish in row Core 3), E1R1 (magenta fish in row Edge 1), E2R1 (yellow fish in row Edge 2), E3R1 (magenta fish in row Edge 3).

## Distinctive spatial patterns between core and edge sampling areas

Figure 3 shows the data obtained from 6 sampling sites, three located at the core of the littoral area (upper rows, light blue) and three located at the edge of the littoral area (lower rows, gray) from 15:20 to 08:00 in 10 100 min-windows. Each tracked individual is shown with a dot, whose distinctive color allows us to follow its position across the 1000 min recording time. In each 100 min-window, the size of the dots represents the time each fish was detected in the tracked position. Transitions between positions are marked with a connecting line. Figure 3 visualizes the day-night spatial patterns of *G. omarorum* in the wild, including individual position and movements, the distinction between residents and visitors, day-night changes, and social interactions. Overall, we were able to track 44 individuals, 31 at the core sampling sites and 13 at the edge sampling sites (Figure 4A).

**Figure 4.**
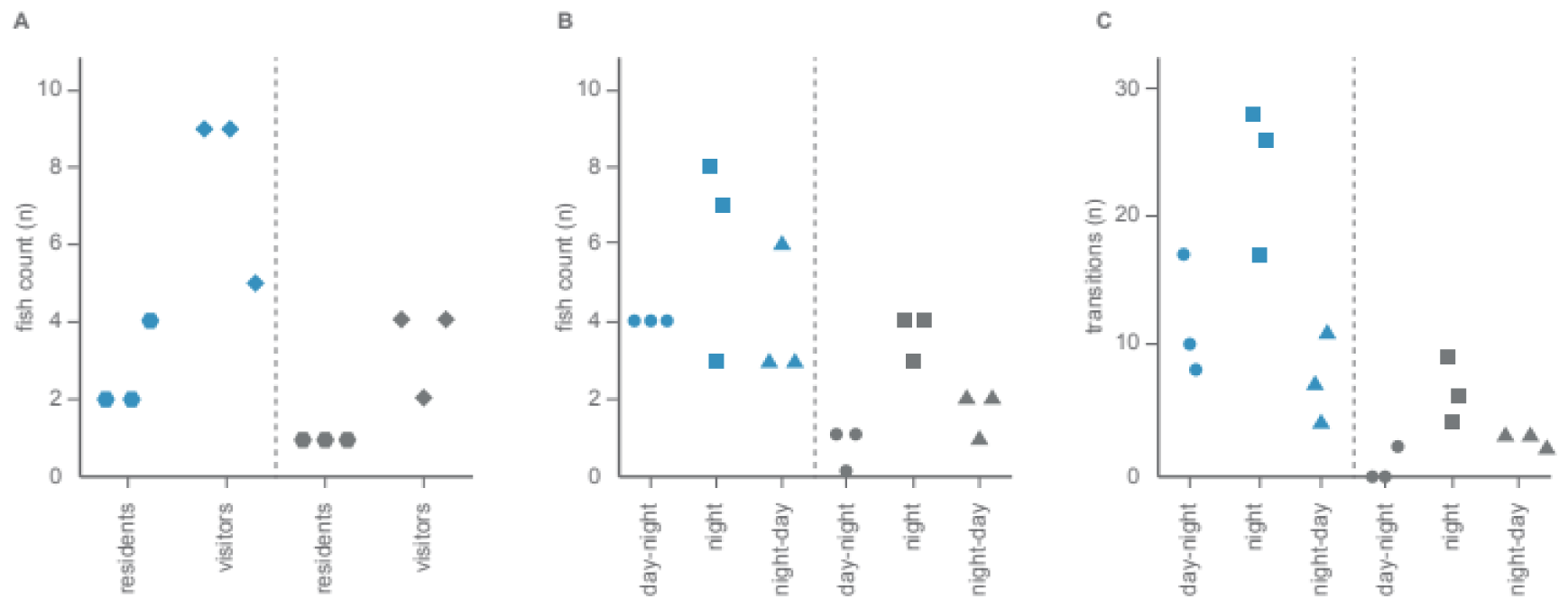
Quantification of fish presence and movements within Core (blue dots) and Edge (gray dots) sampling areas during the 1000 min recording period. Each dot represents the number of fish/transitions found in one sampling site (rows 1-3 in Figure 2). (A) Number of residents (hexagons) and visitors (diamonds) found in the three Core and in the three Edge sampling areas. (B) Number of fish found in the three Core and in the three Edge sampling areas during the night-day (circles), night (squares) and night-day (triangles). (C) Number of fish movements (transitions) found within the three Core and in the three Edge sampling areas during the night-day (circles), night (squares) and night-day (triangles).

We defined as residents those individuals of *G. omarorum* whose presence was identified for more than 60 min either during the first 300 min (day-night transition) or the last 300 min (night-day transition). More residents and visitors were identified in the core sampling areas with respect to the edge sampling areas (Fig. 4A). Only one resident fish was identified in each of the three edge sampling sites (Figure 3). Interestingly, while 1-3 fish were identified as residents of core sampling sites in the day-night transition, only one resident fish was identified in each core sampling site during the night-day transition (Figure 3). Even more interesting is to follow the dynamic transitions of the individuals recorded at each core sampling site in Figure 3. In the upper row, the same two individuals identified as residents at the beginning of the recording period (magenta and cyan dots) remained on site until the end, and adopted the same position within the grid in the morning as they had before sunset. In the second row, one individual was identified as resident of this core sampling site at first (magenta dots), but a second individual that arrived at this sampling patch later remained as the only resident at the end of the night. In the third row, there were 3 resident fish at the beginning of the recording period (magenta, cyan, and yellow dots) but none of them remained until the end of the night.

The nocturnal nature of *G. omarorum* activity becomes evident when observing that during the night, more fish and more individual fish movements were detected in each of the sampling sites (either core or edge, Figure 4B-C). In addition, during the night, more fish and more individual fish movements were detected in core sampling sites than in edge ones (Figure 4B-C). Indeed, during the transitions, especially during the night-day transition, the overall spatial pattern became more similar between core and edge sampling sites with a similar number of identified individuals and a lower number of recorded individual position changes (Figure 4B-C). On several occasions, especially during the night, we were able to identify the simultaneous presence of 2 individuals very close (detected by the same grid electrodes) to each other within the grid system (dots circled in black in Figure 3). Interestingly, whenever these spatial coincidences occurred, one of the fish left the grid afterwards and the one that remained in the grid was always a resident fish.

## The nocturnal increase of EOD rate

Figure 5 shows the mean EOD rate obtained from all the 44 detected individuals (shown with a distinctive color scale in the same way as in Figure 3) in the 6 sampling sites (core sampling areas at the left and edge sampling areas at the right) throughout the day-night-day transitions from the afternoon of the first day (15:20) to the morning of the second day (08:00). The mean EOD rate of resident fish, in which we obtained long-lasting recordings, was interpolated and presented by continuous lines (Figure 5). While the external night extended from 19:00 (sunset) to 07:00 (sunrise) as shown by the solid white-black bar displayed at the bottom of Figure 5, almost no light reached the core sampling areas and only dim light reached the edge ones (gray scale heatmaps in Figure 5). Figure 5 shows that although EOD rate showed a high inter and intraindividual variation during both day and night, there was a consistent nocturnal EOD rate increase, whose onset actually anticipated the beginning of the external night, as well as a consistent EOD decrease anticipating the beginning of the external day. The nocturnal increase of EOD rate always occurred while water temperature was actually decreasing, while the diurnal decrease of EOD rate occurred when the water temperature was at its lowest levels (red-blue heatmaps in Figure 5). Interestingly, as shown in Figure 6, the latencies between sunset and the peak of EOD rate (Figure 6A) were more variable than the latencies between the peak of water temperature and the peak of EOD rate (Figure 6B; Wilcoxon of residuals; p=0,001). These data suggest that the timing of the nocturnal increase of EOD rate was more associated with the daily peak of water temperature than with the timing of the light-dark changes (sunset).

**Figure 5.**
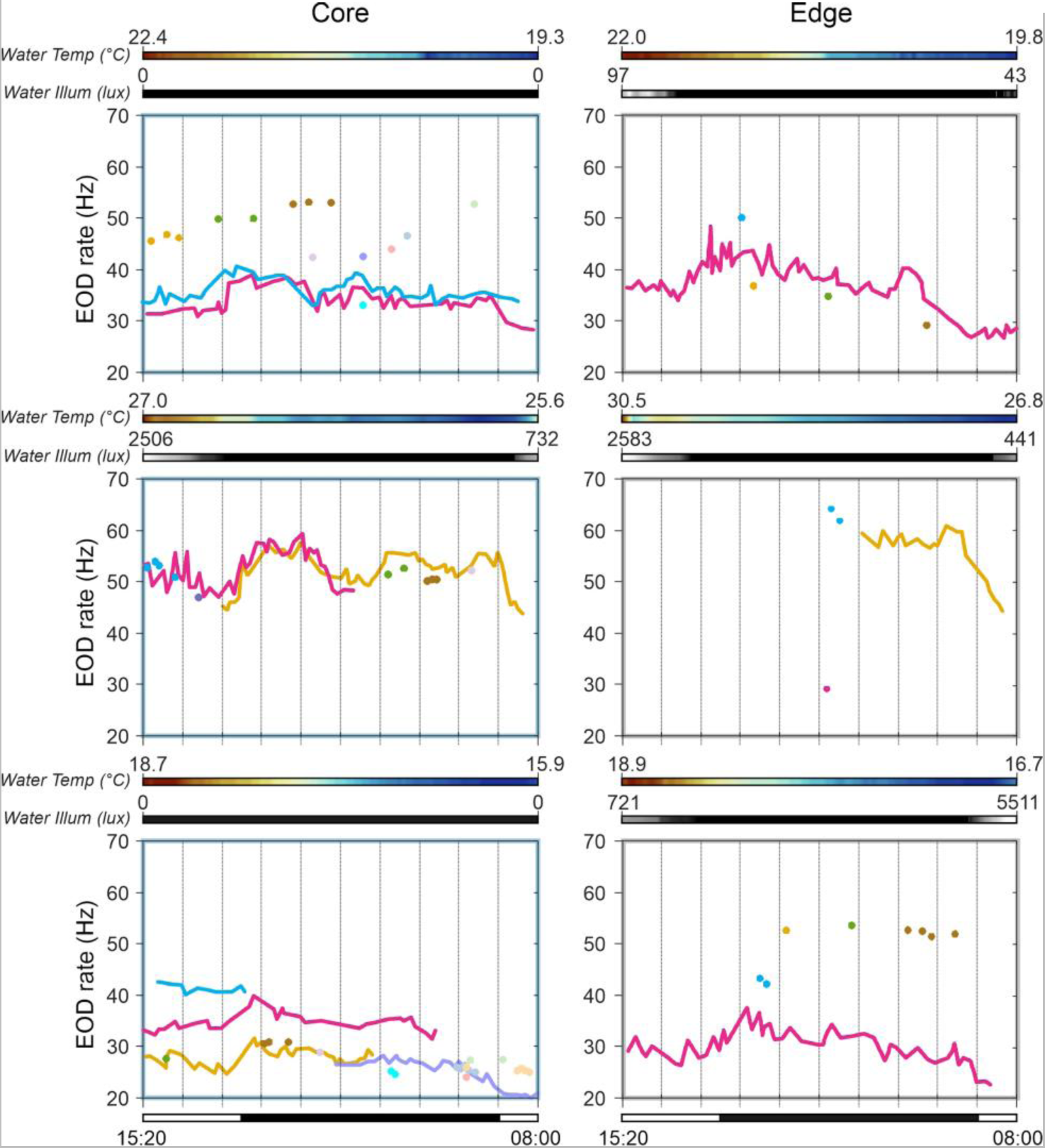
Mean EOD rate across the whole recording period (afternoon of the first day (15:20) to the morning of the second day (08:00)). EOD rates obtained from all detected individuals (n=44) are represented with distinctive colors (same as in Figure 2) in the 6 sampling sites (Core sampling areas at the left and Edge sampling areas at the right). The mean EOD rate of resident fish, based on long-lasting recordings, is interpolated and presented by continuous lines. The mean EOD rate of visitor fish is presented as dots. The solid white-black bar at the bottom indicates the duration of the external night, extending from 19:00 (sunset) to 07:00 (sunrise). Color heatmap shows the water temperature measured at the Core or Edge place. Gray scale heatmaps show the illumination levels measured at the Core or Edge sampling areas. Note that despite the high inter- and intra-individual variations, a consistent increase in nocturnal EOD rate is observed, preceding the onset of the external night, alongside a consistent decrease preceding the external day.

**Figure 6.**
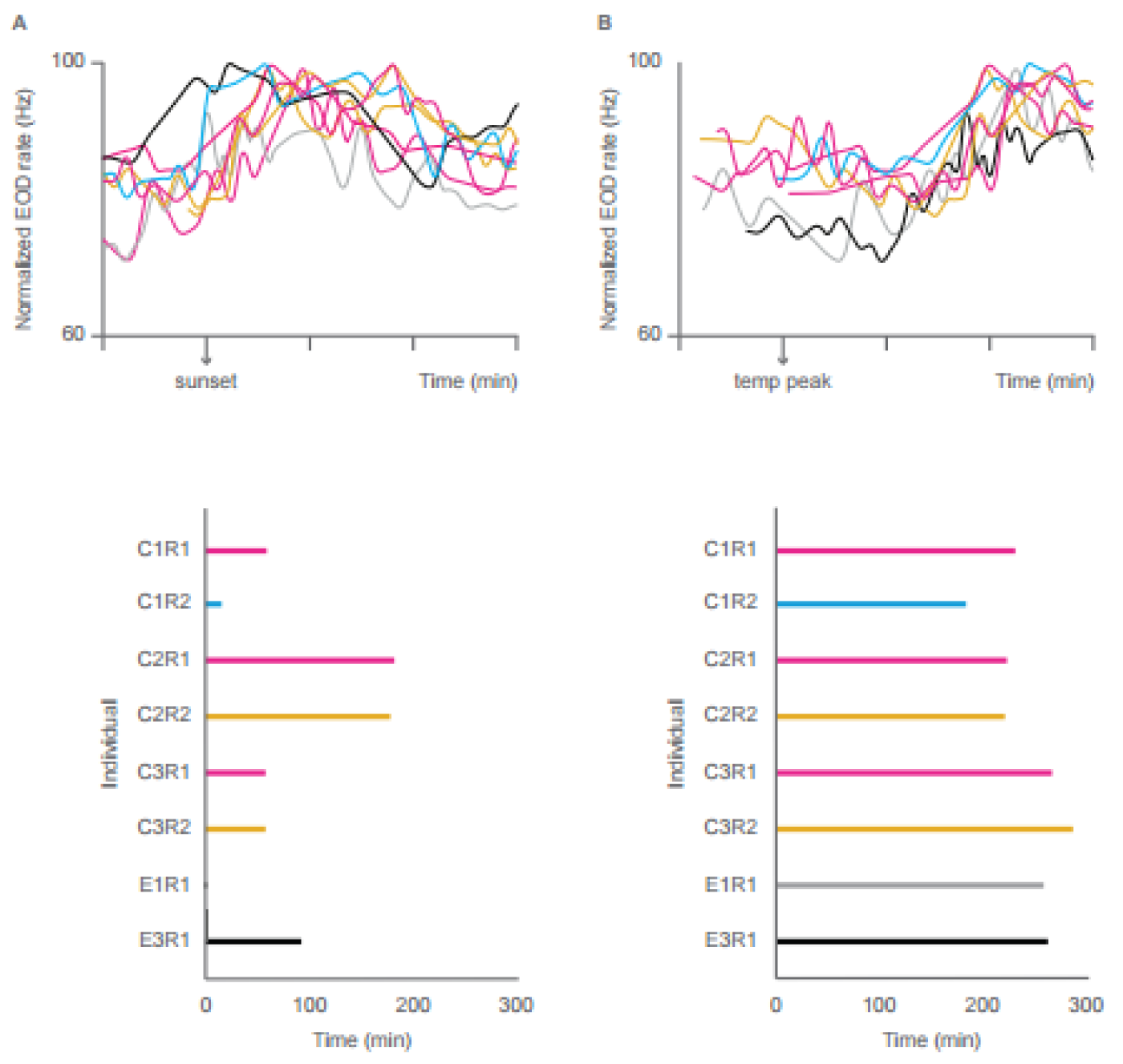
Latency of the nocturnal increase in EOD rate in relation to sunset or to the moment of water temperature peak. (A) Nocturnal increase of EOD rate using sunset (19:00) as time reference. The upper graph shows the normalized mean EOD rate of 8 resident fish from 100 min before sunset to 300 min after sunset. The lower graph shows the time interval (in minutes) from the sunset to the moment of maximum EOD-rate for each resident fish (same individual identification as in Figure 3). (B) The nocturnal increase of EOD rate using the moment of the water temperature peak as time reference. The upper graph shows the normalized mean EOD rate of 8 resident fish from 100 min before the water temperature peak to 300 min after the water temperature peak. The lower graph shows the time interval (in minutes) from the water temperature peak to the moment of maximum EOD-rate for each resident fish (same individual identification as in Figure 3).

## DISCUSSION

The remote tracking of the spatial distribution, locomotor patterns, and behavior of animals in the wild is still challenging for many species due to diverse methodological barriers. In this study, we present the first successful remote tracking of an undisturbed natural population of the pulse-type weakly electric fish, *Gymnotus omarorum,* based on individual-specific traits of their continuously emitting electric signals. Recent previous studies have achieved the individual remote tracking in species of weakly electric fish whose EODs are quasi-sinusoidal by EOD frequency-driven tracking algorithms^3,10,13,14^ and by the development of cutting-edge portable amplifiers^30^. In this study, an updated version of the electric fish loggers together with an adaptation of the tracking algorithms for pulse-fish were important to achieve the individual identification of non-sinusoidal weakly electric fish. Also important to reach this goal was the selection of the proper sampling site and the previous knowledge of the spacing patterns and territorial behavior of the studied species^21,22^. In other words, the interdisciplinary nature of this study allowed us to take advantage of both recently developed electric fish loggers and sophisticated *ad hoc* tracking algorithms to contribute to the understanding of *G. omarorum* daily patterns and territorial behavior. Overall, focusing on continuous recordings during the day-night-day transitions, we were able to follow the location and activity of several individuals *G. omarorum*. Doing so, we also provided evidence on questions long explored in this species but never answered in undisturbed natural populations such as its territoriality, the differential individual spacing within the habitat, and the nocturnal increase in movement, interactions, and electric arousal.

It has been identified for quite some time that the ability of electric fish to generate EODs is an invaluable asset for tracking their location and movements in space. In fact, there have been multiple experimental developments and successful trials in laboratory and natural settings so far^3,10,13–16,31–33^. However, this is the first study reporting remote tracking of an undisturbed population of pulse-type weakly electric fish in the wild. It is interesting to note that this achievement was obtained by the refinement of previously developed recording devices and tracking algorithms rather than by the implementation of novel procedures. For example, both the filtering capacities of the new version of the Teensy microcontroller board and the location of the ground electrode far away from the recording site improved the signal-noise ratio of EOD recordings and therefore their resolution (Figure S1). With respect to the identification of individuals position within the grid recording area, we used an adaptation of the tracking algorithms first developed by Henninger et al^14^, and refined by Raab et al^3^. Although EOD rate is not sinusoidal in pulse weakly electric fish, it is still the main attribute to recognize the identity of individuals *G. omarorum*. EOD waveform and amplitude are too variable to be reliable indicators of fish identity. We thus used EOD rate data to build clusters from the 8 electrodes of the grid system in a similar way as spike sorting from multi-unit electrophysiological recordings^34^. Each cluster was then assigned to one fish, and this was possible partly because of the locomotor patterns of this territorial species, whose individuals tend to move within a small area and to hold the same position for hours. While two simultaneous clusters indicate the presence of two fish underneath the grid system (Figure 2B), merged clusters can also indicate the changes in position of one individual within the grid area across time (Figure 2C).

The continuous remote sensing of individuals *G. omarum* allowed us to confirm some previously known features of the habits and life history of this species and to unravel some new aspects. First, the number of individuals found on each grid patch ranged from 3 to 13 with a maximum of 3 resident fish occurring simultaneously. These findings indicate that the density of this population is actually higher than the previously assessed density of about 1 fish per square meter estimated by diurnal electric census during the non-breeding season^22^. Interestingly, we were able to discriminate between residents and visitors and verify that there were many more visitors than residents. Second, it is clear that more fish and more fish movements were detected during nighttime, which was of course expected for a typical nocturnal weakly electric fish^35^, but has seldom been shown in natural unrestrained populations^14,30^. Third, more fish (residents and visitors) and more fish movements were detected in core sampling sites than in edge ones, which confirms that *G. omarorum* prefers to inhabit the central part of the littoral zone as previously speculated based on daytime manual electrical census^22^. It is remarkable that the spacing patterns of *G. omarorum* and their interactions can be so different between these two microhabitats (core and edge) that are only 2-3 m apart and that do not differ in anything except the proximity of the edge to open water and, consequently, to greater light information. An unexpected result that emerges from our recordings is that fish, especially residents, move only within a small area during the day-night-day transitions. Probably protected and restricted by the dense roots of the aquatic plants, resident fish tend to be detected in the same 1m^2^ area even during their nocturnal active phase. This unexpected stationarity, that was crucial for individual fish identification, is probably typical of the non-reproductive season and therefore may not be observed when the reproductive drive forces adults *G. omarorum* to search for mates.

Since early reports^26^, it has been recognized that the genus *Gymnotus* is highly aggressive and territorial. In particular, the non-breeding aggressive behavior of the study species, *G. omarorum*, has been extensively studied in laboratory and semi-natural settings^20,21,36^, and has become a valuable teleost model system for the understanding of neuropeptidergic and non-gonadal steroidal modulation of aggression^36–40^. In a laboratory arena that mimics the natural conditions of *G. omarorum* habitat, Perrone et al^36^ nicely showed how dyadic agonistic encounters mediate territory acquisition and its defence. After a short highly aggressive contest, the dominant fish keeps the territory, monopolizes the only shelter, and excludes the subordinate from the territory every time it tries to return. Although the remote tracking of *G. omarorum* in the wild did not include video images, our results suggest that agonistic interactions between residents and between residents and visitors mediate territoriality also in natural conditions. For example, in the second row of the core sampling sites in Figure 3, it seems clear that the yellow resident broke into the grid area during the night, interacted with the magenta fish, and excluded it, keeping the territory until the end of the recording period. It is also evident that resident fish excluded intruders (visitors) in almost all (5 out of 6) the recording sites presented in Figure 3. Even more suggestive of agonistic interactions is the observation that every time (8 out of 8) two fish interacted in close vicinity (dots circled in black in Figure 3), one of them retreated from the grid area afterwards while the other remained in the territory as a resident. Finally, site fidelity, the tendency to return to a previously occupied location, has been largely linked to territorial behavior in a wide variety of species^41–44^. As until now individual identification of pulse-type weakly electric fish in the wild had required invasive manoeuvres, site fidelity has been assumed but never confirmed in any territorial species of weakly electric fish. For example, serial electric censuses during successive days on the same location used in this study have indicated the presence of individuals *G. omarorum* on the same spot (L. Zubizarreta, personal communication). However, this procedure cannot determine that each time a fish is detected, it is effectively the same animal. The results presented in the first row of the core sampling sites in Figure 3 contribute a very robust confirmation of the site fidelity of *G. omarorum*. The magenta and cyan fishes that moved within the grid sample area quite a lot during nighttime, preferred the same spot of the grid to rest before and after the nocturnal active phase. Indeed, this precise site fidelity was actually observed in all the resident fish that remained within the grid sampling area during the whole recording period.

Day-night changes in electric behavior have been well-documented in several species of weakly electric fish in laboratory settings at constant temperature^26–29,45^. As a common trait across species, these day-night changes include a nocturnal increase in EOD rate, which has been interpreted as a nocturnal electric arousal that improves electrolocation and electrocommunication resolution during the active phase. Interestingly, this nocturnal electric arousal persists in motionless fish and under constant darkness in the wild while water temperature is actually decreasing^25,45,46^. A previous study carried out in the same core sampling site we used in this study showed that restrained individuals of *G. omarorum* exhibit a robust nocturnal EOD rate increase^25^. This nocturnal electric arousal occurred without changes in water illuminance (i.e., in constant darkness) but was likely synchronized by daily variations of both water temperature and social cues^25^. In this study, we obtained EOD rate data from the 44 detected freely moving individuals of *G. omarorum*.

Interestingly, visitors (individual dots in Figure 5) tended to have higher EOD rates than residents, probably because visitors were more actively exploring around the grid sampling area than residents. On the other hand, the long lasting continuous recordings of several resident fish allowed us to observe, for the first time in an undisturbed natural population, a clear EOD rate increase during the night, whose amplitude was similar to that previously reported in restrained individuals of this species^25,29^ and also similar among fish recorded from core or edge sampling sites. Both the increase and decrease of EOD rate anticipated sunset and sunrise, respectively. But, while we captured the peak of the nocturnal increase of EOD rate during the night, we cannot be certain about the timing of its minimum because the EOD rate was at its lowest values at the end of the recording period.

Temperature is a major environmental factor, particularly relevant for ectotherms, because of its direct effects on metabolic processes. For example, when *G. omarorum* is subjected to short term gradual temperature changes, the higher the temperature the higher the EOD rate^47^. However, the nocturnal increase in EOD rate occurs after water temperature has reached its maximum, and coincides with the declining phase of the temperature cycle (Figures 5 and 6). This opposite response to changes in water temperature suggests that temperature is acting in at least two different ways, as the well-known determinant factor in metabolism, and as a synchronizing signal from the environment. In line with previous reports^25^, we show that the onset of the nocturnal increase of EOD rate in undisturbed individuals was loosely associated with the beginning of the night but tended to be more synchronized with the peak of the daily changes in water temperature. Temperature, as an environmental timing cue, has been less explored than the light-dark cycle, despite being particularly relevant in the context of climate change. Among many other implications, global warming is leading to an attenuation of temperature cycles and an increased uncoupling of the photoperiod from the thermoperiod^48^.

The relevance of this study lies in its inherently interdisciplinary approach. On one hand, the refinement of cutting-edge recordings and data processing procedures were key to provide a new successful example of undisturbed tracking of natural populations of aquatic animals based on individual physiological cues. On the other hand, the evidence of individual day-night spatial patterns provided by this study confirmed relevant information about spacing, habits, and behavior of this species previously suggested by non-automated procedures. On top of this, the inclusion of different environmental cycles as determinants of natural behavior broadens the scope of this study, allowing an ecological perspective. Behavioral studies of ectotherms in the natural habitat provide an excellent opportunity for assessing the contribution of temperature as an environmental cue, which is of uttermost importance to assess the impact of global warming in natural populations.

## LIMITATIONS OF THE STUDY

We recognize several limitations of this study, most of them derived from the development of innovative methodologies in undisturbed natural settings. For example, the ground electrode was located in the water only 50-100 cm apart from the grid sampling area. Thus, if one fish remained close to the ground electrode, it would have been detected in all the 8 recording channels and act as a confounding factor. Second, the identification of daily spatial patterns would be better if we had obtained longer recordings. On the other hand, several operational decisions made during data processing can be considered arbitrary such as the categorization of residents or the time delay of 10 min for a fish that leaves the grid sampling area to be considered as a different fish if an EOD of similar rate was detected. Finally, body size, which is the major determinant of the agonistic outcome in this species, could not be estimated by the tracking algorithms because the position of the animal is much more impactful on the amplitude of EOD recordings than body size.

## ACKNOWLEDGMENTS

This research was partially funded by CSIC, Universidad de la República, Uruguay, Programa de Grupos I+D – 2018 # 92 to AS and by DFG grant 501755140 to JB.

## AUTHOR CONTRIBUTIONS

Conceptualization, A.S., J.B., A.M., F.P.; Methodology, J.B., A.M., F.P., S.M.; Software, J.B., F.P.; Formal Analysis, F.P., A.M.; Investigation, A.S., J.B., A.M., F.P.; Resources, A.S., A.M.; Writing – Original Draft, A.S., A.M.; Writing – Review & Editing, A.S., J.B., A.M., F.P., S.M.; Supervision, A.S., J.B.; Funding Acquisition, J.B., A.S.

## DECLARATION OF INTERESTS

The authors declare no competing interests.

## EXPERIMENTAL MODEL AND SUBJECT DETAILS

Fieldwork was performed in Laguna de los Cisnes, Uruguay (205 ha, 34° 480 S, 55° 180 W), which consists of a three-part interconnected shallow system of freshwater (maximum depth 5 m) with no inputs of salt or brackish water (Figure 1A,^22^). The experiments were conducted during the non-breeding season at the peri-equinox period (March 10-12, 2023 and October 10-12, 2023) under a natural light-dark cycle of 12:12. Periodic light and temperature measures were taken each 30 min, inside the water close to the recording sampling sites and outside the water (HOBO-MicroDAQ: UA-002-08). Measurements range: temperature, −20° to 70°C (−4° to 158°F); light, 0 to 320,000 lux (0 to 30,000 lumens/ft2). The study species, *Gymnotus omarorum* ^1*7*^, is the sole species of weakly electric fish present in this study area^18^. The littoral area of the lake is blanketed by a dense strip (5–40 m width) of free-floating aquatic macrophytes that cover the sampling area (Figure 1B), with *G. omarorum* typically resting among the roots of these plants. Data was collected from an unperturbed natural population. As illustrated in Figure 1B, two grid systems were placed on top of the natural floating vegetation, one closer to the shore (aka Core, light blue) and another in the outermost limit of the natural vegetation patch (aka Edge, gray). Each grid system consisted of an array of eight electrodes distributed on a 1m^2^ surface separated by 40 cm in positions that maximized the spatial resolution that results from the combined analysis of the eight simultaneous recordings (Figure 1C). Consequently, this grid array assured the detection of the EODs of any individual of *G. omarorum* in any position within the recording area. Due to the nocturnal habitats of *G. omarorum,* recordings (from 15:20 to 08:00) focused on the night and the day-to-night and night-to-day transitions during three nights. As shown in Figure 3, this recording period was divided in 3 epochs: 1) the day-night transition (first 300 minutes); 2) the night involving 400 minutes of recording; and 3) the night-day transition (last 300 minutes). General protocols of *G. omarorum* handling in the wild were approved by the institutional Ethical Committee for the use of experimental animals (Facultad de Ciencias, Universidad de la República, CEUA-1304), but this study did not actually involve animal collection nor manipulation.

## METHOD DETAILS

Electrical recordings were made using eight carbon rod electrodes (50 cm long) which were placed below the plant coverage that reaches 30 cm under the water surface. In approximately the same water depth, *G. omarorum* is usually found resting among the roots^25^. Data were recorded over three nights (3/10/2023, 10/10/2023, 10/11/2023). Two recording systems were used. All edge recordings and the 3/10/2023 core recordings were filtered and differentially amplified against a reference electrode in 50cm distance (30×gain, 100 Hz high-pass filter, 7 kHz low-pass, Teensy-Amp R1.0, https://github.com/janscience/Teensy_Amp/tree/main/R1.0), digitized with 12bits at 20kHz by a Teensy 3.5 microcontroller (www.pjrc.com), and stored in continuous 1 minute wave files on µSD cards for offline processing. The 10/10/2023 and 10/11/2023 core recordings used the 8-channel Teensy_Amp R4.0 (https://github.com/janscience/Teensy_Amp/tree/main/R4.0, <u>jlm Innovation</u>, Tübingen, Germany) with 10x inverting preamplifiers and a separate analog-digital-converter (two PCM1865, Texas Instruments) that digitized the signals with 16bit at 48kHz controlled by a Teensy 4.1 microcontroller (www.pjrc.com) with additional 8MB PSRAM chip for buffering the data. The Teensy_Amp R4.0 input signals are connected via 100kΩ to common ground and thus do not require a separate reference. Logger software for both systems is based on the TeeRec library (https://github.com/janscience/TeeRec) and the “8-channels-logger” and “R40-logger” sketches of the TeeGrid project (https://github.com/janscience/TeeGrid). The custom-built recording systems were powered by phone power bank batteries (5 V, 10-20 Ah). Recordings were analyzed with Matlab routines for EOD detection and EOD rate calculation.

The key differences between the two recording systems are whether an external reference electrode is needed (R1.0) and whether the onboard ADC of the microcontroller (R1.0) or a separate ADC chip (R4.0) is used. In supplemental Figure S1 recordings of the two systems are compared. In the Laguna 50 Hz hum is quite strong and harmonics of 50 Hz extend over the whole spectrum. Finding the right position of the reference electrode of the R1.0 system is crucial. In the wrong position the hum gets too strong and a proper detection of EODs gets impossible. This is quite some effort, since for each position of the reference electrode the recorded wave files have to be taken from the SD card and analysed on a laptop, before a decision can be made about the best position. The R4.0 system avoids the need of a reference electrode by tying the floating signals via 100 kOhm resistances to the internal battery ground. This ground effectively follows the average signal over the 8 channels and this way removes most of the external noise. In addition, the ADC of the Teensy microcontroller used by the R1.0 system effectively uses 12bit and is influenced by voltage changes induced by writes to the SD card. The separate ADC used by the R4.0 system internally uses 24bit, does not suffer from SD writes and has under ideal conditions a much lower noise floor. In addition, a single PCB of R1.0 amplifies 2 channels. We used 4 of the R1.0 PCBs to assemble an 8-channel amplifier. A single PCB of the R4.0 system amplifies and digitises 8 channels. The new R4.0 system thus provides a compact and easy to use solution for multi-channel recordings of EODs.

### Algorithms for the estimation of fish position

The presence of individuals *G. omarorum* relied on detecting EODs in one or more of the electrodes distributed in the grid array (Figure 2A). The voltage traces in each electrode were bandpass-filtered (Butterworth filter, third order, 300-2000Hz). From this filtered data the peak-to-peak amplitude *A_i_* of the detected brief EOD pulse was obtained for all 8 electrodes. EOD amplitude was then used to estimate the location of each fish following Henninger et al^14^ as shown in Figure 2B. The fish position was estimated from the amplitudes *A_i_* at the *n=*8 electrodes *i* at position *e_i_* as a weighted spatial average, given by:

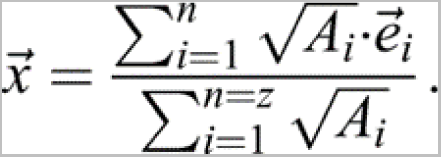

The EOD of a fish can cause a very large amplitude on nearby electrodes because of the electric field’s reciprocal dependence on distance. This effect results in a relatively large localization error, if a simple weighted spatial average is used, because the position estimate is pulled towards the strongest electrode. The localization error is reduced by using the square root of the EOD amplitude, √Ai, as a weight, which reduces the impact of electrodes with large EOD amplitudes (Henninger et al., 2020).

### Algorithms for individual identification

Individual fish were identified based on a cluster analysis and merged over time according to their EOD rate and position between analyzed files (Figure 2C). The primary assumption for EOD detection was that the EODs of pulse-type weakly electric fish are brief signals with almost no temporal coincidence. Using the x and y coordinates of the positions obtained as described in the previous section, the maximum number of clusters to consider (max) was selected. This number was then used to obtain the within-cluster sum of squares (WCSS) for different values of k=1:max using k-means. The elbow point of WCSS was determined as the value where the maximum of the second derivative was found. Subsequently, a new k-means analysis was conducted with k being the previously obtained value. For each cluster the EOD rate was calculated.

If this fish moves, the EOD amplitude detected by each electrode will change and eventually a change in the position of the fish will be detected, this in turn will generate a new cluster. To account for this drift in signal a post-analysis was performed to merge contiguous clusters with similar EOD rates (<10% difference, connecting lines between dots in Figure 3). Clusters in between files were merged using both spatial and EOD rate criteria. Only clusters with at least 100 EODs will be recognized as one individual of *G. omarorum* located at a certain position within the grid (dots in Figure 2). When the identified fish leaves the grid recording area, EODs will no longer be detected and the fish will be considered gone, unless a fish with the same EOD rate is detected again in less than 10 minutes. Given the previously reported inter-individual distance and the speculated high site fidelity of *G. omarorum*^22^, this approach proves to be the most feasible for individual identification.

### Fish classification

Individuals *G. omarorum* detected within the grid system were categorized as residents or visitors. We recognized as residents those fish whose presence was identified for more than 60 min either during the first 300 min (day-night transition) or the last 300 min (night-day transition). We recognized as visitors those fish whose presence was identified for shorter periods throughout.

## QUANTIFICATION AND STATISTICAL ANALYSIS

### EOD rate and indexes

EOD rate was calculated as the inverse of the inter EOD interval. A mean EOD rate was calculated for each 1 minute file and interpolated. Due to the interindividual differences in EOD basal rate, changes in rate were normalized to allow comparisons among individuals. The nocturnal increase index (NI) was calculated as follows:

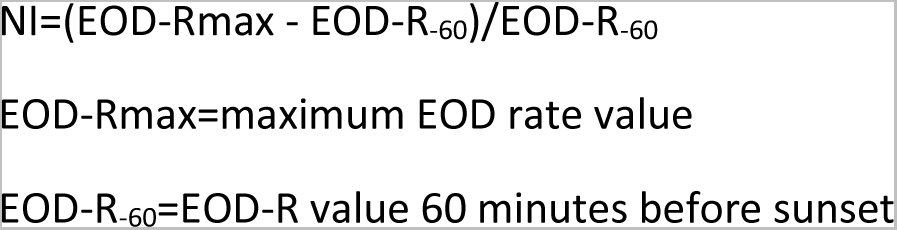

To evaluate the association of the nocturnal increase of EOD rate with environmental variables, we calculated the latency in minutes between the sunset and the peak of EOD rate and the latency in minutes between the peak of water temperature and the peak of EOD rate.

Statistical procedures were carried out using the software PAST^49^. Time associations between light and temperature cues with the nocturnal increase in EOD rate and the diurnal decrease in EOD rate were compared using the nonparametric Wilcoxon signed ranked test. Values of p < 0.05 were considered statistically significant.

## SUPPLEMENTARY MATERIAL

**Figure S1.**
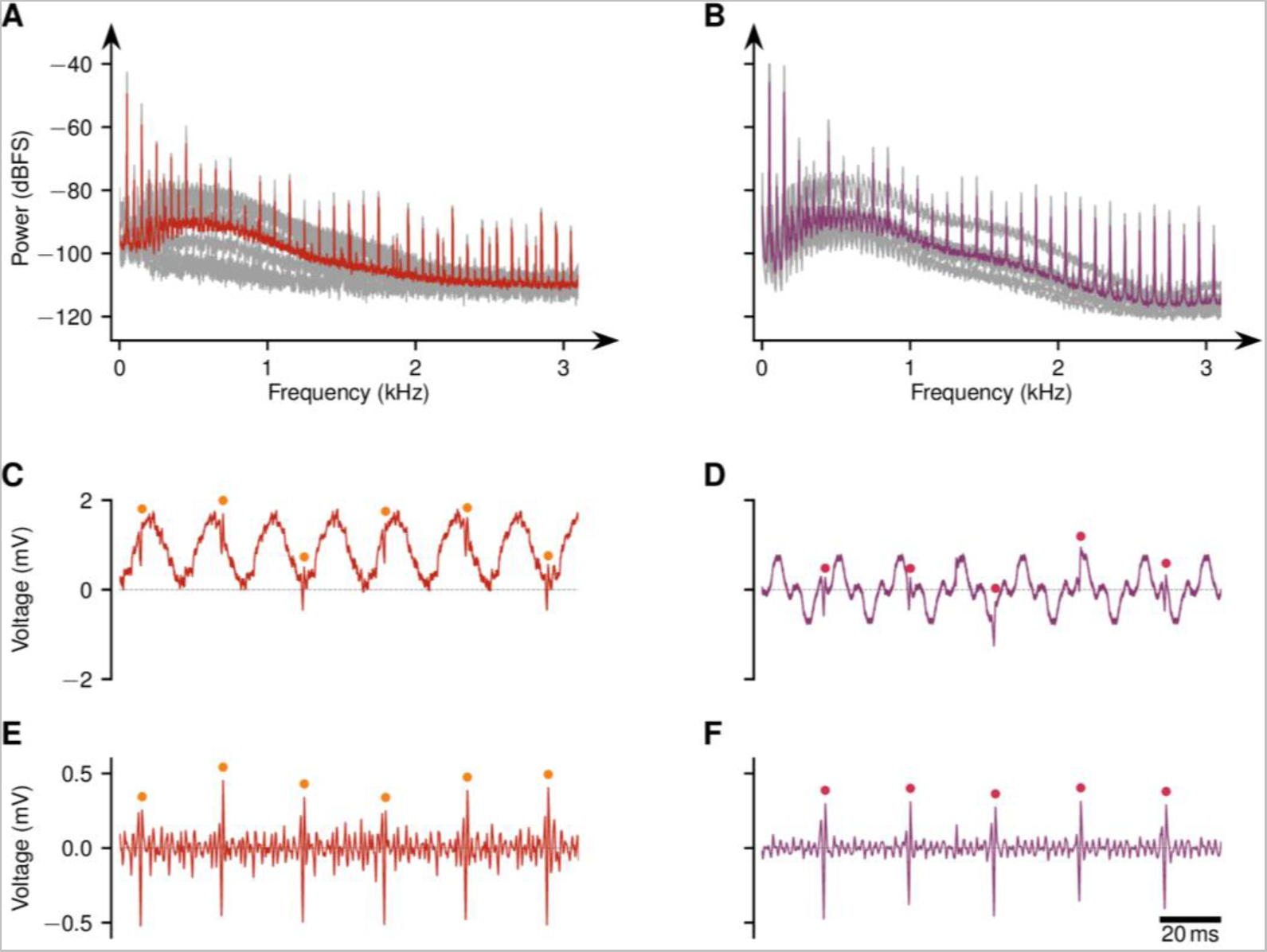
Comparison of the Teensy_Amp R1.0 (left) and the Teensy_Amp R4.0 8-channel loggers. (A, B) Power spectral densities of a 15 s long recording (each channel grey, average over channels in color) demonstrate the strong 50 Hz power hum (peaks) present in the Laguna. The overall shape of the power spectra varies a lot between channels and over time and is rather similar between the two amplifier systems. (C, D) The 50 Hz hum clearly dominates over the EODs (marked by dots) in the 150 ms long example traces. Again, the exact waveform of the noise varies over channels and time. Despite its highpass filter, the Teensy Amp R1.0 has a small offset bias (here about 1 mV). (E, F) After band-pass filtering between 300 Hz and 2 kHz the hum is suppressed and the EODs clearly stand out (dots). The newer Teensy_Amp R4.0 produces a better shaped noise floor that despite a similar rms power is less peaky. Also note that the examples for the Teensy_Amp R1.0 show recordings with the reference electrode placed to result in the lowest possible noise level. In contrast, the Teensy_Amp R4.0 produces recordings of similar if not better quality without a reference electrode that needs to be positioned.

## Notes

### Competing Interest Statement

The authors have declared no competing interest.

