## Supplementary figures and images for "Tracking spatial patterns and nocturnal arousal in an undisturbed natural population of the pulse-type weakly electric fish *Gymnotus omarorum*"

### Supplementary figure 1

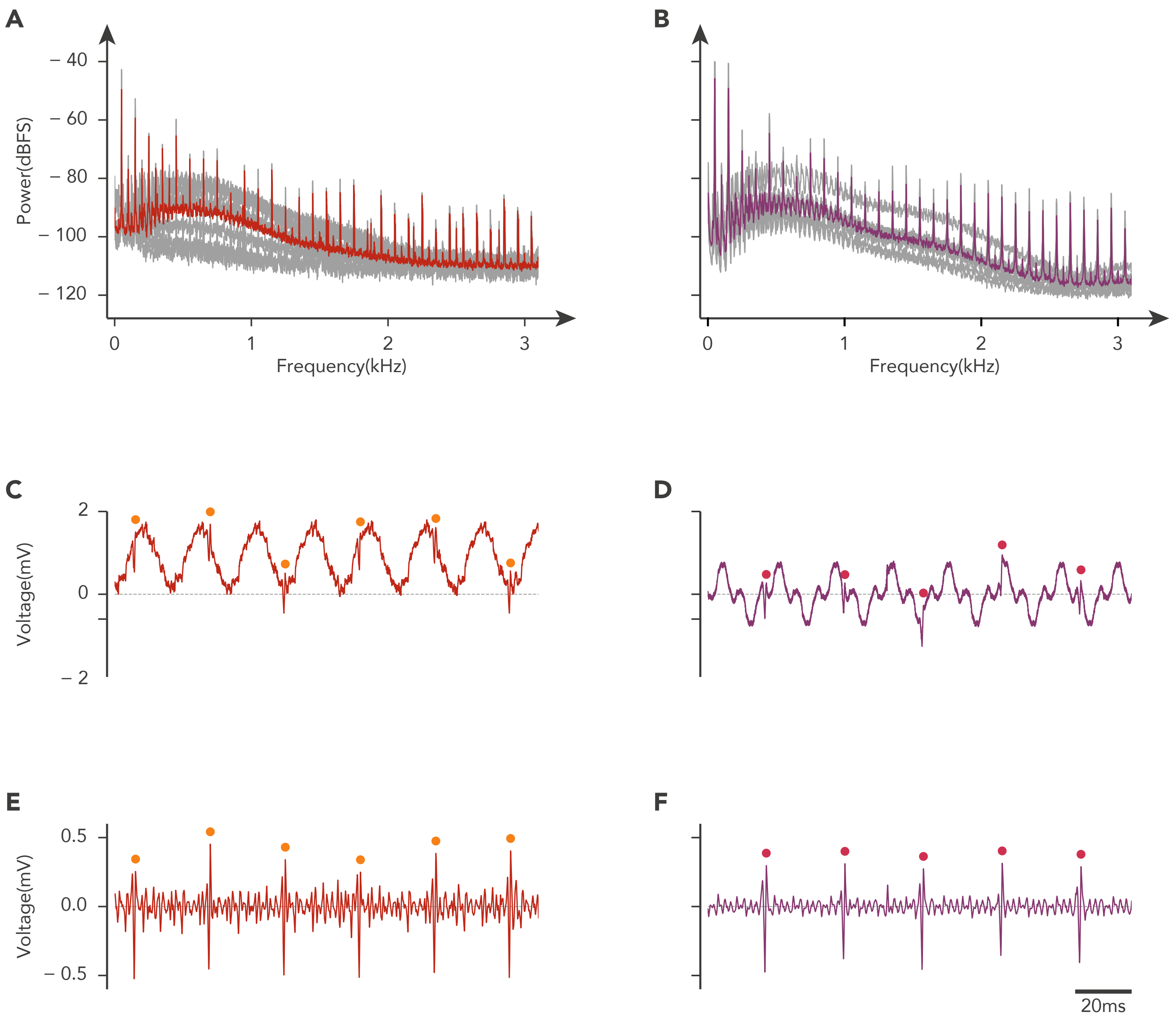
